# Beyond linearity: Quantification of the mean for linear CRNs in a random environment

**DOI:** 10.1101/2022.08.26.505415

**Authors:** Mark Sinzger-D’Angelo, Sofia Startceva, Heinz Koeppl

## Abstract

Molecular reactions within a cell are inherently stochastic, and cells often differ in morphological properties or interact with a heterogeneous environment. Consequently, cell populations exhibit heterogeneity both due to these intrinsic and extrinsic causes. Although state-of-the-art studies that focus on dissecting this heterogeneity use single-cell measurements, the bulk data that shows only the mean expression levels is still in routine use. The fingerprint of the heterogeneity is present also in bulk data, despite being hidden from direct measurement. In particular, this heterogeneity can affect the mean expression levels via bimolecular interactions with low-abundant environment species. We make this statement rigorous for the class of linear reaction systems that are embedded in a discrete state Markov environment. The analytic expression that we provide for the stationary mean depends on the reaction rate constants of the linear subsystem, as well as the generator and stationary distribution of the Markov environment. We demonstrate the effect of the environment on the stationary mean. Namely, we show how the heterogeneous case deviates from the quasi-steady state (Q.SS) case when the embedded system is fast compared to the environment.

## 1 Introduction

Regulation of gene expression is generally well-modelled by stochastic chemical reaction networks (CRNs) and can be successfully represented using other approaches only under certain conditions [1]. The modelled system, e.g., a single gene or a small gene network, exists in the context of a larger gene network and the cellular environment (Fig. 1a). It is crucial to account for the influence of this context onto the modelled system while not inflating the model complexity. A standard approach for this is to represent all external influence on a given system as a consolidated stochastic process [2, 3]. In the case when the environment affects the rates of the studied process, we must consider a CRN with modulated reaction rates.

**Fig. 1.**
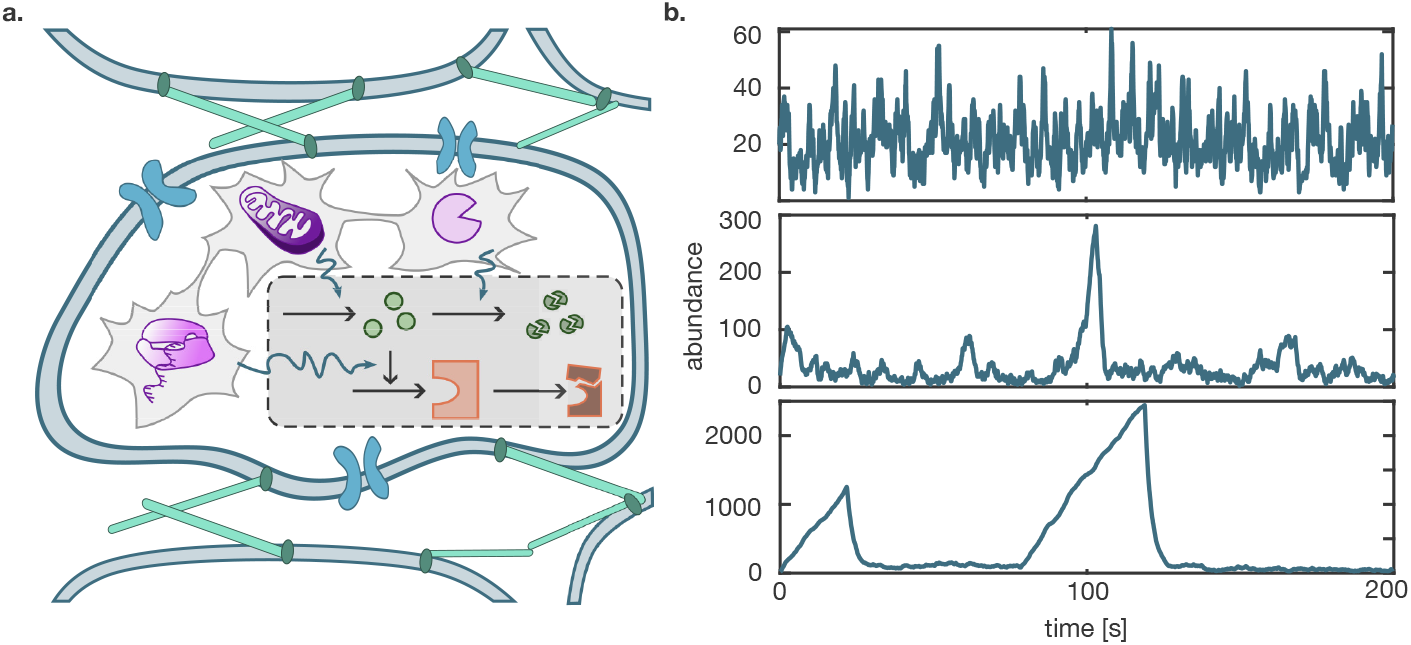
**a.** The cartoon shows an example subsystem of a reaction cascade that is embedded into a cellular environment that includes RNA polymerase activity, ATP availability, and the concentration of a degradation enzyme. **b.** Trajectories of a birth-death-process with heterogeneous death rate *μ*, modelled as a birth-death process on different time scales. On the fast time scale, the trajectory resembles an unmodulated birth-death process (upper panel). On the intermediate time scale, triangular excursions appear while *μ* = 0 (middle panel). On the slow time scale, the excursions become more pronounced (lower panel). The mean death rate is the same for all cases.

In modelling studies, protein degradation, or the death process, is often assumed to be an indiscriminate, constant-rate process. This, however, is known to not always be the case. Namely, protein degradation is regulated via a ubiquitin-proteasome pathway, which allows fine-tuning the degradation rates of specific proteins [4–6]. Further, controlling targeted protein degradation is a promising route for treating various health conditions such as cancer, neurodegenerative diseases, metabolic disorders, infections, and inflammatory diseases [7]. Previously, it was not possible to perturb the protein death process in a controlled way, which hindered the detailed modeling of these processes. However, recently, new techniques for this were introduced [7, 8]. Thus, it is now relevant and feasible to develop new modelling strategies that would incorporate modulated protein degradation rates.

The study of general chemical reaction networks dates back to the 60s [9], after [10], [11], [12] and others had pioneered small case studies. Typical quantities of interest are the probability distribution or the generating function and moments both in the transient and stationary behaviour. [13] still emphasized the analogy of the mean equations with the deterministic counterpart. However, the following works [9, 14] demonstrated that bimolecular reactions cause a characteristic deviation of stochastic systems when the system is not in the thermodynamic limit, highlighting the need for other techniques. Early works computed the generating function and mean at stationarity for small bimolecular networks comprising two balanced reactions [15]. The observation that bimolecular reactions result in an unclosed mean equation brought more attention to unclosed (hierarchical) moment equations. Unclosed moment equations have prompted researchers to pursue several strategies. Among them, stochastic simulations for systems with highly abundant components are time-consuming and therefore computationally prohibitive. Equally prohibitive is the use of the master equation, which gave rise to more efficient hybrid methods [16–18]. Overall, previous studies developed and extensively explored various moment closure schemes [19–22], identified the limitation of moment closures with regards to bimodal distributions [18] and extinction [23], and pointed out the limited local character of moment closures [24]. More recently, linear programming under positive semi-definite constraints was employed to compute upper and lower moment bounds [25–27] and approximated the moments in the case of tight bounds.

For linear reaction networks, the first- and second order moment dynamics are well-known both in the transient and stationary phase [28]. In particular, the stationary mean of the stochastic system equals the equilibrium concentration of the corresponding deterministic system [29]. We have a clear understanding of the case when a linear subsystem is modulated in its zeroth-order reactions. Several research groups characterized the second-order moments [28, 30, 31] for this case and [3] introduced a method with which they (approximately) simulated marginal subsystem trajectories for various example networks. Zeroth-order modulation by a linear environment is included in the theory on linear systems, because the joint subsystem-environment network is linear. Non-linear reaction systems were embedded in a linear environment while assuming linear subsystem-environment interaction [32], i.e., zeroth-order modulation (in both directions).

The effect of a random environment on the noise in a subsystem, i.e., on the variance, is well-studied. Many research groups adopted the decomposition of noise into intrinsic and extrinsic components since its introduction in the seminal paper [33], and this concept was expanded upon in later studies [30, 34–36]. In contrast, we investigate the effect of the random environment on the mean of a subsystem species. This effect is visible when first-order reactions are modulated as the following simple example illustrates. Consider a birthdeath process with birth rate λ and death rate *μ* whose mean approaches λ/*μ* in the equilibrium. Zero-order modulation corresponds to a modulation in the birth rate, i.e., stochastic λ with mean 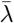. By the linearity of the function λ ↦ λ/*μ* and the linearity of the mean, the heterogeneous mean 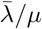 does not reflect the heterogeneity at the mean level. However, if *μ* is stochastic, the non-linearity of *μ* ↦ λ/*μ* invalidates this argument. Instead, the excursions in the abundance that emerge during *μ* = 0 can distort the heterogeneous mean away from 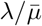 (Fig. 1b). In the experimental context, this means that even bulk data carries the fingerprint of a heterogeneous environment.

The simple birth-death process in a random environment has been studied in queuing theory. It corresponds to an *M/M/*∞ queue, i.e., a queuing system with infinitely many servers and exponential arrival and serving times. The birth rate is the rate of arrival, while the death rate corresponds to the service rate. For a Markov environment, the stationary distribution, stationary factorial moments, and the transient evolution equation of the moments were derived in [37]. Bimodal stationary distribution was reported. Complementary to this result, the special case of the random telegraph modulated service rates was fully characterized in [38]. It was extended to simultaneous birth- and death-modulation by a semi-Markov environment in [39].

In this work, we consider the general class of linear reaction systems (subsystems) whose reaction rates are modulated by a discrete state Markov environment. As our main contribution, we analytically express the stationary mean for this class of CRNs in terms of the subsystem rate constants, the environment generator matrix and the environment equilibrium distribution. This extends the existing results in queuing theory to more reaction channels than the birth-death channel. Furthermore, we propose a new method that quantifies the shares that the environment states contribute to the stationary mean of a subsystem species. Finally, we analyze the deviation from the Q.SS behaviour in a variety of case studies to illustrate several non-linear effects of the random environment.

## 2 Model class

### 2.1 General CRN model

A CRN is defined as a system of *M* chemical reactions R_1_,…, R_*M*_ involving *d* reactant species **X** = [*X*_1_,…, *X_d_*] with the stoichiometric substrate and product coefficients *S_ij_* and *P_ij_*, respectively, and reaction rates *c_j_*, where *i* ∈ {1,…, *d*}, *j* ∈ {1,…, *M*}:

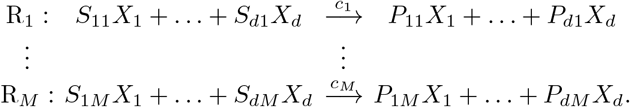

The propensities of the system transitioning between states follow the massaction kinetics, with the propensities 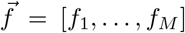 being a function of the stoichiometric constants and the reaction rates, where the function form depends on the values of the stoichiometric constants. Note that reactions of higher than the second order, i.e., reactions with 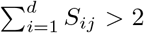, are generally not considered, as the probability of such reactions occurring is negligibly low compared to the reactions of the lower orders.

### 2.2 Homogeneous linear CRN

Consider a *linear* CRN with the vector 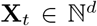 of species counts, namely, a CRN where all reactions follow mass-action kinetics of zeroth or first order [28]. By this, we mean that the vector of propensities for its *M* reactions is of the form

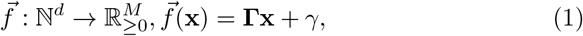

where **Γx** and *γ* are the vectors with the propensities of the first-order and the zero-order reactions, respectively. Next, we construct the stoichiometric matrix that stores the change vectors for each reaction:

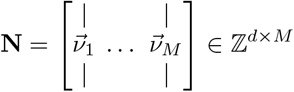

via *N_ij_* = *P_ij_* – *S_ij_*. This allows to write down the evolution of the mean:

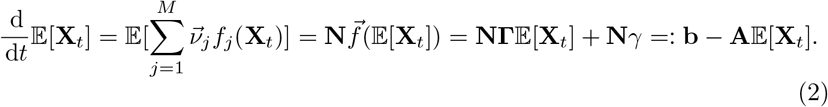

The matrix **A** = –**NΓ** captures the first-order reaction rates, while the vector **b** = **N***γ* accounts for the zero-order reaction rates. Throughout this paper, we assume that in the homogeneous case **A** has only eigenvalues with positive real part.

### 2.3 Linear CRN in a random environment

We have just described the homogeneous case. Next, we embed the linear chemical reaction network in a random environment. We assume that *Z* is a stationary continuous-time Markov chain (CTMC) on a discrete state space 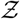. The environment modulates **X** by *Z*-dependent propensities which replace Eq. (1) by 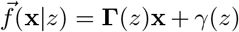 with corresponding family of matrices **A**(*z*) and vectors **b**(*z*) as defined in Eq. (2). We refer to such a linear CRN in a random environment also as a heterogeneous, or modulated, linear CRN. Throughout this paper we assume that **A**(*z*) has only eigenvalues with non-negative real part for each 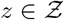. In contrast to the homogeneous case, 0 may be included as an eigenvalue.

Most prominently, **Z** can be a CRN on the state space of (environmental) species counts 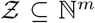 [3, 40]. In this case, we call **X** the *subsystem* of the joint reaction network (**X**, **Z**). The equation that governs the mean now reads

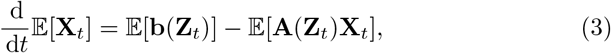

involving non-linear modulation terms 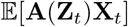.

### 2.4 Quasi-steady state reference model

With a quasi-steady state (Q.SS) assumption on the environment, we aim to map a heterogeneous linear CRN to a homogeneous one. To this end, we replace the subsystem propensities 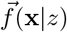 by 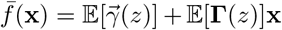. Even though the quasi-steady state method has seen applications with much less coarse approximations in stochastic models, i.e., at the level of the master equation [41], we use the term in the described way at the mean level. The corresponding first-order rate matrix and zero-order rate vector, see section 2.2, are 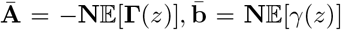. In the case of 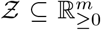 and if **Γ** or *γ* depend linearly on **z**, then 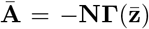 or 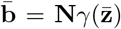, respectively. The mean Eq. (3) reads

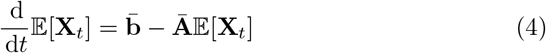

and converges to the equilibrium

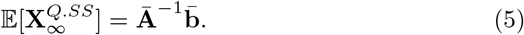

We use the Q.SS model as a reference model. The deviation of the heterogeneous from the homogeneous reference model quantifies the effect of the random environment on the subsystem. Quasi-steady state model reduction is typically justified by the assumption that the environment operates on a faster time scale than the subsystem. This particular rather coarse Q.SS model cannot be justified as a model reduction technique, when taking into consideration that more elaborated and superior techniques [41–43] were developed. However, we justify our Q.SS in the reverse perspective: In the modeling context, when we build a hierarchy of models, we may start with homogeneous reaction rates, i.e., a deterministic constant environment. As we pass to heterogeneous rates, the rate means are kept constant, e.g., if the environment means were obtained from separate measurements.

Then the heterogeneous case deviates from the homogeneous Q.SS case. Is it necessary to include a higher-order moment analysis in order to detect the deviation? We demonstrate how the fingerprint of the heterogeneous rates emerges already in the stationary mean of the subsystem.

As a point of departure, we first portray a base case without deviation at the mean level, i.e., the stationary mean of the Q.SS model coincides with the heterogeneous stationary mean. Suppose, the environment modulates the subsystem only via zero-order reactions, i.e. **A**(*z*) ≡ **Ā** = **A** is independent of the environment, while **b**(*z*) may maintain dependencies. Then Eq. (3) reduces to

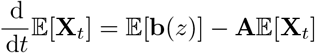

with equilibrium state 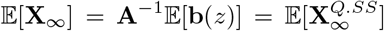. Only in the second (and higher) order moments the exact network deviates from the Q.SS model [30]. The additional term which enters in the variance expressions is commonly interpreted as extrinsic noise, opposed to the intrinsic fluctuations that are attributed to the subsystem alone [36]. While the stationary mean is not affected by the zero-order modulation, this is much different when we allow modulation of first-order reactions. The question how the random environment affects the subsystem even on the mean level guides the remainder of this paper.

## 3 Results

We analytically express the stationary mean for the class of CRNs introduced in the previous section. The expressions only depend on the subsystem rate constants and standard characteristics of the Markov environment that we specify as follows. Denote by 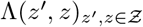 its generator and by *π*(*z*) its stationary distribution, i.e., **Λ***π* = 0. Introduce the notation Λ_0_(*z*):= –Λ(*z*, *z*) = ∑_*z*′≠*z*_ Λ(*z*′, *z*) for the total exit rate. Let 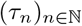 be the jump times of *Z*. These induce the discrete-time Markov chain (*W_n_*)_*n*_:= (*Z*(*τ_n_*))_*n*_ with transition kernel

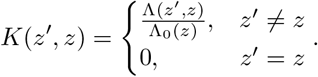

The process *W_n_* is called embedded discrete-time Markov chain that corresponds to the jump epochs of *Z*(*t*) [44]. Define

### Proposition 1

*Let W be the embedded chain of a Markov jump process on* 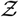 *with stationary distribution π*(*z*) *and total exit rates* Λ_0_(*z*), 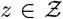. *Define* 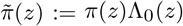. *Then* 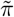 *satisfies the stationarity condition for the embedded chain, i.e., is an unnormalized stationary distribution of W*.

*Proof* For the check of the stationarity condition, see Appendix A.1.

### 3.1 Auxiliary machinery

For the computation of the stationary mean, we use the average values of the process **X** at the end of intervals [*τ_n_*, *τ*_*n*+1_] in environmental state *z*. The computation of these average values is facilitated by the fact, that conditional on the environment the subsystem expectation progresses linearly.

#### Definition 1

Define 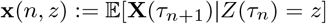 and 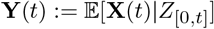.

Conditional on the history of *Z*, the subsystem **X** is linear and hence **Y** evolves like

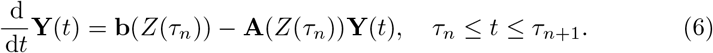

We note that the value *Z*(*τ_n_*) is the only information in the history of the environment that governs the evolution (locally in time). For the value **Y**(*t*) global in time, the initial jump value **Y**(*τ_n_*) is needed, which depends in a more complex way on past values of *Z*. This is solved interval-wise by

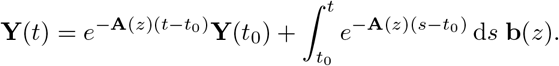

Choosing *t*_0_ = *τ_n_* and *t* = *τ*_*n*+1_, we just expressed

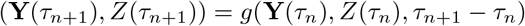

for a deterministic update function *g*. By the update characterization of Markov chains, the independent waiting times (*τ*_*n*+1_ – *τ_n_*)_*n*_ and independence of *τ*_*n*+1_ – *τ_n_* from (**Y**(*τ_n_*), *Z*(*τ_n_*)), we get that the pair (**Y**(*τ_n_*), *Z*(*τ_n_*))_*n*_ forms a Markov chain. Furthermore, by the tower property of conditional expectations, we obtain the following link between **x** and **Y**

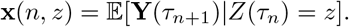

At stationarity, **x**(*n*, *z*) does not depend on *n* anymore, because (**Y**(*τ*_*n*+1_), *Z*(*τ_n_*)) has the same distribution as (**Y**(*τ_n_*), *Z*(*τ*_*n*–1_)). Hence, we can drop *n*. More formally:

#### Definition 2

Define 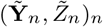 as the stationary version of the Markov chain (**Y**(*τ_n_*), *Z*(*τ_n_*))_*n*_ and 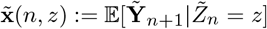. Define 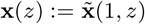.

#### Proposition 2

*The* **x**(*z*), *defined in definition 2, satisfy the linear equations*

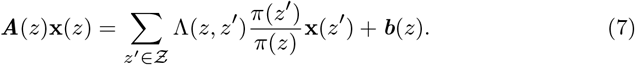

*Suppose that π*(*z*) *satisfies detailed balance, then*

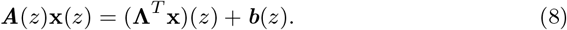

*Proof* See Appendix A.2.

*Remark 1* Define **y**(*z*):= *π*(*z*)**x**(*z*) and denote by **y**^*T*^ = [**y**(*z*_0_)^*T*^, **y**(*z*_1_)^*T*^,…] the concatenated vector for an enumeration of the environment state. Further define the block matrices

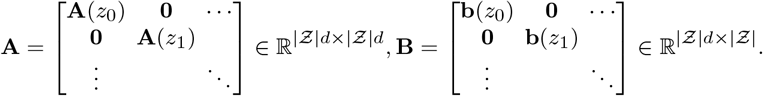

Then Eq. (7) can be written as

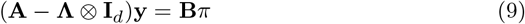

where **I**_*d*_ is the (*d* × *d*) identity matrix, ⊗ is the Kronecker- or tensor-product and 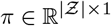 is the stationary probability vector. Denote by **x**^*T*^ = [**x**(*z*_0_)^*T*^, **x**(*z*_1_)^*T*^,…] the concatenated vector for an enumeration of the environment state. Furthermore consider **b** = **Be** with 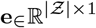 the vector with ones in all entries. Then Eq. (8) can be written as

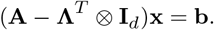

### 3.2 Exact stationary mean evaluation (ESME)

We computed the average values of the process **X** at the end of intervals [*τ_n_*, *τ*_*n*+1_] with environmental ‘label’ *z*. With these auxiliary quantities, we obtain the following main result for the stationary mean.

#### Theorem 3

(ESME) *Let **X** be a linear CRN in a random environment as described in section 2.3. With* **x** *defined as in definition (2) and π the stationary distribution of Z, it holds*

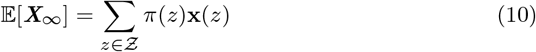

*Proof* See next section and Appendix A.2.

*Remark 2* If we invest equation (9), the resulting equation (10) for the ESME is rewritten in matrix-vector notation

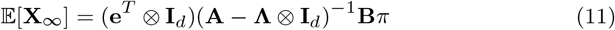

with ⊗ the Kronecker-product, **I**_*d*_ the *d* × *d*-identity matrix and 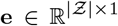 the vector with ones in all entries. For *d* = 1, the expression reduces to

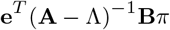

for diagonal matrices **A**, **B**. This is in agreement with the expression obtained in [37, theorem 3.1]. If *π*(*z*) satisfies detailed balance, then it also holds

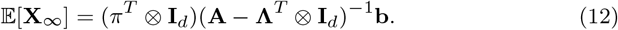

### 3.3 Proof of the ESME expression

In this section, we provide the essential part of the proof for the main theorem 3 (ESME) in detail. This equips the reader with the intuition on quantifying the shares that environmental states contribute to the stationary mean. We define these in the next section. Note that our proof deviates from those in queuing theory in that it does not pursue a generating function approach. For technical parts of the proof we refer to Appendix A.2.

By the ergodic theorem, we can move from the ensemble mean to the temporal mean

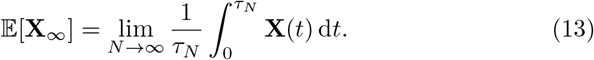

Next, we partition the time axis at the jump times *τ_n_* both in the numerator and denominator to obtain

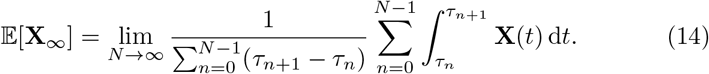

The summands (in both sums) are ordered by their appearance in time. The idea is now to sort them by the values of *Z*(*τ_n_*). This is achieved by multiplying each summand with unity of the form 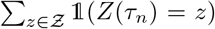 and changing the order of summation. The outer sum is now indexed by 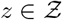. The next idea is to normalize the numerator and denominator by *N* to get empirical averages. As *N* → ∞, the ergodic theorem allows us to move back to expectations. In the denominator, this reveals the average waiting time as a mixture of inverse exit rates:

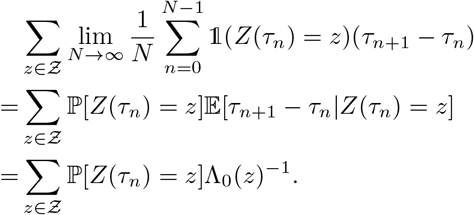

In the numerator, the contributions of *Z*(*τ*n**) = *z* to the sum is handled analogously by moving from the empirical mean to the expectation

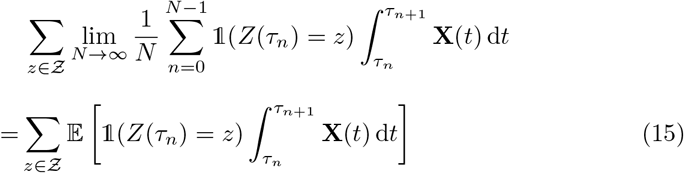

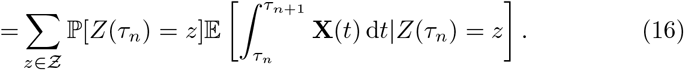

The integrals in Eq. (16) evaluate to

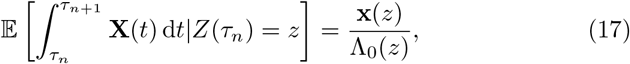

which is computed in the Appendix A.2 along with the remaining calculations.

*Remark 3* The expression (10) has a rather astonishing interpretation that can be understood as a consequence of the waiting time paradox [45]. Suppose the system operates in stationarity. For simplicity, we assume *X* is one-dimensional. If we choose a random time *t*, then with probability *π*(*z*) we hit an interval *τ_n_* ≤ *t* < *τ*_*n*+1_ with a ‘label’ *Z*(*t*) = *z*. Call this event *I_z_*. The time point adds 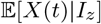 to the mean if we think of the stationary mean as 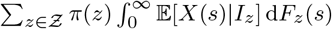, and *F_z_*(*s*) is the distribution of *s* = *t* – *τ_n_* when in *Z*(*t*) = *z*. Looking at the structure of expression (10), the integral evaluates to 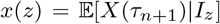. Why is this paradox? We might ad hoc assume that *t* lands, on average, at some centered location within the interval [*τ_n_*, *τ*_*n*+1_], in particular it is, on average, smaller than *τ*_*n*+1_. The progression of 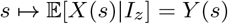, see Eq. (6), is strictly monotone. Imagine it is increasing (decreasing). Then *t* < *τ*_*n*+1_ implies 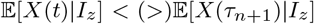. By the monotonicity of the expectation, this implies 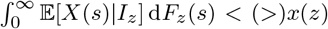. In contrast, Theorem 3 informs us that equality holds. This can be understood as a consequence of the waiting time paradox. A uniformly random time point *t* satisfies that *τ*_*n*+1_ – *t* is exponentially distributed with parameter Λ_0_(*z*) by the memory-less property of the exponential distribution. However, due to symmetry reasons in the uniform choice of *t*, the distance *t* – *τ_n_* of the last *Z*-jump in backwards time is also exponentially distributed with parameter Λ_0_(*z*). To deviate from the main line of thought, the interval that *t* lands in has, on average, twice the expected length. This paradox is resolved by the size bias effect: longer intervals have a higher chance to be hit by the point *t*. The explicit size-biased distribution of *τ*_*n*+1_ – *τ_n_* is provided in [45]. Returning to the main line of thought, a randomly chosen time point is actually an average endpoint, regarded from the perspective of a randomly chosen interval. The difference lies in the random choice of a time point versus the random choice of an interval. The waiting time paradox permits the slim formulation of Theorem 3, once the recursion in Proposition 2 is solved.

### 3.4 Quantification of environmental shares

The stationary mean is a composite result of the subsystem existing in different environmental states. Thus, we aim to quantify the share that each environmental state contributes to the value of the stationary mean. First, let us fix a subsystem species 1 ≤ *i* ≤ *d* and consider its stationary mean 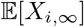. When we computed the stationary mean following Eq. (14), we sorted the summands by the environmental states *Z*(*τ_n_*) = *z*. Inspired by Eq. (15), we define the *environmental share* that environmental state *z* contributes to the stationary mean of species *i*, as

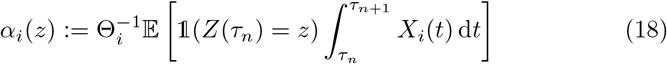

with the normalization

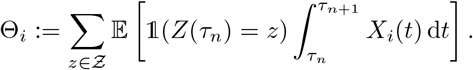

We note that, by definition, 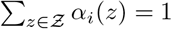. It holds

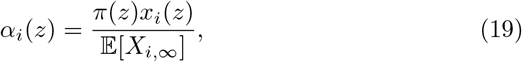

because 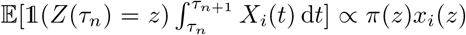 by Eq. (17) and Proposition 1.

## 4 Case studies

We analyze several small chemical reaction networks and gene regulatory motifs, including an example relevant in synthetic biology, to demonstrate how the stationary mean (ESME, Theorem 3) and the environmental shares can provide insight into the properties of the system. ESME requires the generator and the stationary distribution of the Markov environment, as well as the reaction rate constants of the linear subsystem. The generator and reaction rate constants are specified by the model, whereas the stationary distribution can be obtained numerically, or in special cases analytically, e.g., for the class of monomolecular CRNs [46].

### 4.1 Birth-death process in a random environment

First, we illustrate the effect of three different stochastic environments E1 - E3 (Fig. 2a-c) on the stationary mean of a birth-death process *X*. The modulated first-order death reaction can be seen as a bimolecular reaction. Here, we demonstrate the effect of different relative speeds between the environment *Z*_2_ and the subspecies *X*. The environments E1 - E3 modulate the birth and death rates independently. The birth modulation via a birth-death process *Z*_1_ is the same in all cases, while the complexity of the death rate modulation *Z*_2_ increases.

**Fig. 2.**
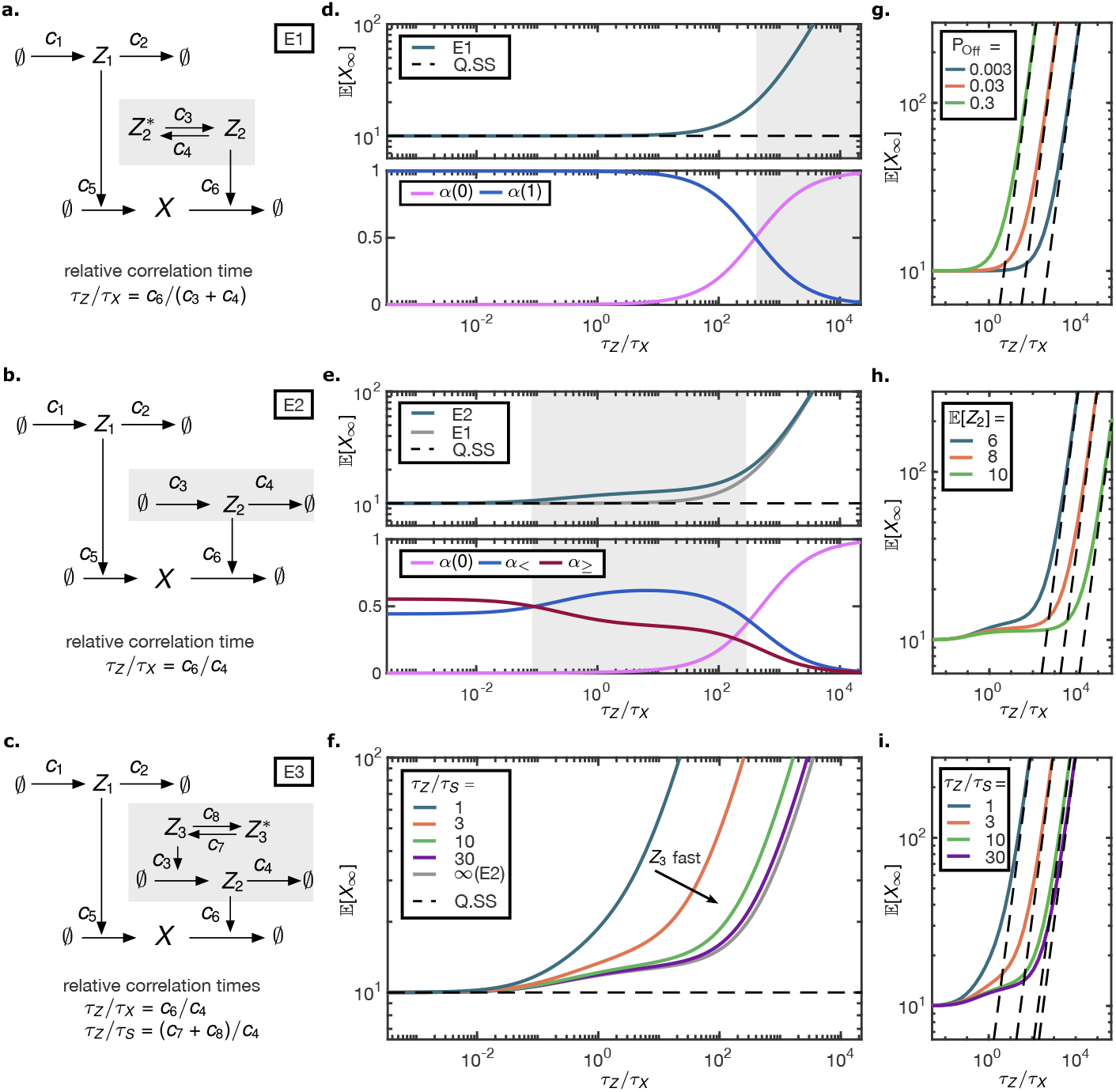
**a,b,c** Environment schemes for the birth-death process *X*. The two processes *Z*_1_ and *Z*_2_ modulate the birth and the death rate, respectively. **d** *Upper panel*, **e** *Upper panel*, **f** The Stationary mean as a function of the relative correlation time systematically exceeds the Q.SS mean, to which it converges for fast environment (valid Q.SS assumption). **d** *Lower panel*. The contributions of *Z*_2_ to the stationary mean (E1). For slow environment the share of *Z*_2_ = 0 dominates. **e** *Upper panel*. Three regimes for E2 can be distinguished: the asymptotic, the intermediate and the degenerate regime. The matched plot of E1, Eq. (23), shows agreement in the fast and slow regimes and a discrepancy for the intermediate regime. *Lower panel*. The contributions of *Z*_2_ = *z* for *z* ∈ {0}, 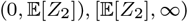) to the stationary mean indicate the distinct regimes. **f** Mutable *Z*_2_ synthesis (E3): The different curves reflect different relative speeds between environmental components *Z*_2_,*Z*_3_. For fast *Z*_3_ the muting is neglectable and the environmental effect E2 from **e** is recovered. For slow Z3 the deviation from the Q.SS mean increases. **g,h,i** The asymptotic behaviour for E1 - E3 as the relative subsystem speed tends to ∞. The log-log plot demonstrates that 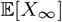 increases proportional with the relative correlation time. The proportionality constant was computed according to section 4.1.4 and is shown as dashed line. Different values for the **g** percentage of inactivity 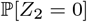 (E1), **h** environment mean (E2), **i** relative speed between environmental components (E3) are shown. The relative correlation times *τ_Z_*/*τ_X_* = *c*_6_/(*c*_3_ + *c*_4_) (E1), *c*_6_/*c*_4_ (E2, E3) progress from slow to fast subsystem (hence slow environment) in increasing direction. All parameter choices kept the Q.SS mean 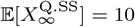 and the ratio *c*_5_/*c*_6_ = 1 fixed. For E2, E3 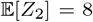 was chosen unless indicated otherwise, while in E1 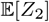 was calibrated to match 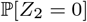 with E2. For E3 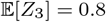 =0.8 was held constant when varying *τ_Z_*/*τ_S_* = (*c*_7_ + *c*_8_)/*c*_4_.

#### 4.1.1 Death modulation via random telegraph (E1)

First, we considered the scenario where the death rate is modulated by a two-state Markov process (Fig. 2a). A two-state modulation highlights the effect that the Off state 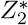 has on *X*. When the instant decay rate is zero, the molecular numbers of *X* increase unboundedly. We call these phases excursions. We expect the stationary mean to depend on (i) the length and (ii) the frequency of excursions, because (i) the temporal average during one excursion increases with the length of the excursion, and (ii) if excursions occur more frequently, the excursion average value is weighted more strongly. The frequency can be characterized by 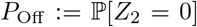, whereas the length of excursions is proportional to the relative correlation time. The autocorrelation function of the random telegraph process, as well as the birth-death process, is of the form *e*^−*t/τ*^, where *t* is the time lag and *τ* is the correlation time. Thus, for the random telegraph process *Z*_2_ with On and Off switching rate *c*_3_, *c*_4_, the correlation time is *τ_Z_* = (*c*_3_ + *c*_4_)^−1^, while for the birth-death process with constant degradation *c*_6_ it is 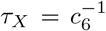. With these definitions, the expression of the stationary mean (derived using ESME, Eq. (12), see Appendix B.1) takes the form

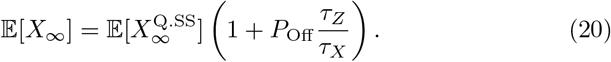

In the queuing literature, [38] already derived an equivalent expression in terms of rate constants, along with other characteristics for the model, e.g., stationary distribution, using a generating function approach. The bracket term is larger than 1, yielding a systematic positive deviation compared to the Q.SS as a reference model (Section 2.4). Note that this can be expected from the moment equations. The mean equation for *X* is

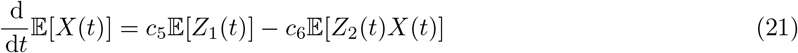

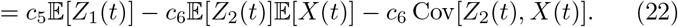

The first two terms in Eq. (22) would yield the Q.SS dynamics. Since *Z*_2_ and *X* are negatively correlated, the stationary mean is larger compared to the Q.SS mean. As Eq. (20) shows, the deviation is proportional to the separation of time scales *τ_Z_/τ_X_* and to the probability 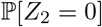.

Figure 2d (upper panel) portrays the stationary mean 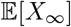 as a function of the relative correlation time *τ_Z_/τ_X_*, given that the Q.SS mean and *P*_Off_ are constant. Increasing the relative correlation time can be achieved by either accelerating *X* or decelerating *Z*_2_. Figure 2d (lower panel) shows how the share of *Z*_2_ = 0 increases with the mean waiting time in *Z*_2_ = 0 (or decreases with increasing speed of *Z*_2_). In the asymptotic regime *τ_Z_/τ_X_* → 0, the stationary mean reaches the Q.SS mean, confirming that the Q.SS assumption is valid for sufficiently fast *Z*_2_ or, equivalently, slow *X*.

#### 4.1.2 Death modulation via birth-death process (E2)

We next investigated whether the generic expression Eq. (20) still holds when the death modulator *Z*_2_ is itself a birth-death process (Fig. 2b). To this end, we altered 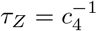 and 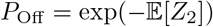 and asked whether

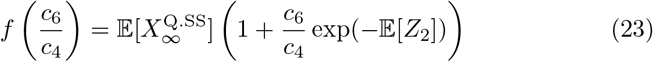

approximates 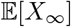, computed via ESME in Appendix B.2. The approximation is valid for the extreme cases of (i) *c*_4_ large compared to *c*_6_ or (ii) *c*_4_ small compared to *c*_6_ (Fig. 2e, upper panel). Namely, we found 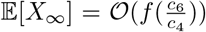 for (i) 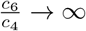 and (ii) 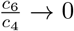. The means 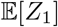 and 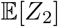, as well as *c*_5_/*c*_6_, were fixed.

Second, we partitioned the states of *Z*_2_ into three classes: the zero state, the non-zero states below the mean, and the states equal to or above the mean. The relative speed *τ_Z_/τ_X_* of the environment defines which of these three classes dominates in terms of the corresponding environmental share, i.e., the effect on the stationary mean. In order to interpret the deviation of E2 from E1, we quantified environmental shares according to the three classes of environment states:

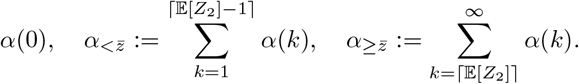

We found that, depending on the relative speed of the environment, the subsystem can be in one of three phases (Fig. 2e, lower panel). For a small *c*_6_/*c*_4_ ratio, the non-zero shares 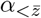 and 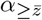 both contribute significantly, with 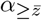 showing a slight dominance over 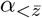, while the share of *α*(0) is negligible. For a medium *c*_6_/*c*_4_ ratio, 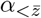 takes the lead in dominance, while *α*(0) is still negligible. Finally, for large *c*_6_/*c*_4_, the share *α*(0) dominates the contribution to the mean. Returning to the upper panel of the Figure 2e, we confirm that the mean 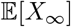 as a function of *τ_Z_/τ_X_* undergoes the same phase transitions.

The qualitative behavior for large *τ_Z_/τ_X_* is driven by unbounded excursions in the state *Z*_2_ = 0. However, in biological systems, these are generally prevented by, e.g., a leakage in the death rate. Upon introducing a base death rate λ_0_, we expect the stationary mean to saturate at the upper bound as the relative environmental speed approaches zero. To demonstrate this, we generalize the propensity of the death reaction to *f*(*x|z*) = (*c*_6_*z* + λ_0_)*x*.The figure B1 indicates four qualitatively different regimes of the stationary mean over the relative correlation time. In particular, the three phases of the model E2 analysis (Fig. 2e.) persist, whereas the fourth phase with the largest *τ_Z_/τ_X_* reaches saturation.

#### 4.1.3 Mutable synthesis of the modulator (E3)

We next considered the case where the modulating process *Z*_2_ is itself modulated by another process, *Z*_3_ (Fig. 2c). The modulator 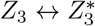 is a two-state Markov process that acts as a switch with On (*Z*_3_) and Off 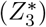 states, switching rates *c*_7_, *c*_8_, and correlation time *τ_S_* = (*c*_7_ + *c*_8_)^−1^. Here, *Z*_2_ can be seen as a regulatory protein produced from a promoter *Z*_3_ that alternates between On and Off states. Then *Z*_2_ is a Markov-modulated birth-death process and, as such, represents an example of a non-Markovian death rate modulator. Since the joint environment (*Z*_2_, *Z*_3_) is Markovian, ESME applies (see Appendix B.3).

We varied the relative correlation time 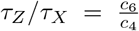 while keeping the ratios 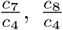, and 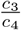 constant. Furthermore, we varied the relative speed of the environmental components by additionally varying the relative correlation time 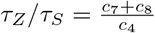 while keeping the fraction of time the modulator is active, 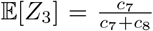, constant. For a large relative speed 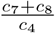, the original model with a constant birth rate 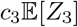 is recovered as expected.

Comparing the stationary means at different relative switching speeds of the modulator (Fig. 2f), we found the following. For slower relative speeds, the deviation from the Q.SS mean becomes more pronounced already at smaller correlation times 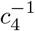, and the intermediate phase vanishes. Meanwhile, Figure B2a visualizes how the entry into the degenerate regime depends on *τ_Z_/τ_S_* and 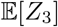. The muting prolongs the excursions of *X* in the zero or sub-average *Z*_2_ states.

#### 4.1.4 Stationary mean for slow environment

We considered each of the environments E1 - E3 without leakage in the death rate of *X* and analyzed the behaviour for a slow environment and a fast subsystem, i.e., *τ_Z_/τ_X_* → ∞. We aimed at isolating the effect of the time scale separation. For this purpose, we kept the means and the relative speed of the environment components, as well as *c*_5_/*c*_6_, fixed. As the plots of the environmental shares (Fig. 2d,e, B2b) suggest, the state *Z*_2_ = 0 is the only one that contributes in the limit case *τ_Z_/τ_X_* → ∞. From this, we derived

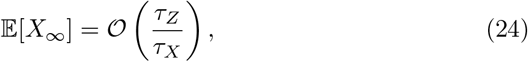

see Appendix B.4, i.e., the stationary mean grows proportionally to the time scale differences. The parallel asymptotes in the log-log plots with unit slope in the Figures 2g,h,i reflect this dependence.

### 4.2 Synthetic controller mitigating the heterogeneous degradation rate

One goal of synthetic biology is to design circuits that are robust to environmental changes. The setpoint objective specifies a target copy number or concentration at which the species of interest should be kept robustly, i.e., the concentration is supposed to re-adapt when environmental changes perturb it. We considered the setting in which an environmental birth-death process modulates the degradation of *X* (Fig. 3a). A controller species senses the environment and acts on the birth rate of *X* to attenuate the effect and achieve the setpoint objective for *X* [47, Fig S.9c]. We chose the Q.SS mean as the setpoint, and as the deviation measure we employed the relative deviation

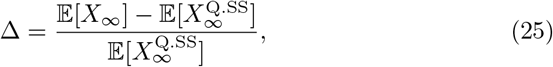

or the accuracy measure Δ^−1^, respectively. In the previous sections we saw that *Z* = 0 can be the main driver of deviations from the Q.SS. The controller *U* works against this effect: during phases of otherwise unbounded *X* excursions, the birth rate of *X* is now down-regulated by *U* and, in the extreme case, comes to a halt at a plateau (for details see Appendix B.5).

With stochastically independent birth and death modulation that we considered so far, the stationary mean was not affected by fluctuations in the birth modulation, and we could apply the Q.SS assumption on *Z*_1_. Here, on the contrary, the time scale of the controller species *U* matters in reacting robustly to the environment. Figure 3b depicts the stationary mean of *X* for different controller speeds *c*_2_/*c*_6_. The deviation Δ gets more pronounced for a slower controller, whereas a fast controller achieves better accuracy Δ^−1^. For the effect to become apparent, the environment needs to be sufficiently slow (*τ_Z_/τ_X_* large). When the controller operates slowly, it does not have the attenuating effect. In this regime, the target species achieves a base accuracy that depends on a given environment speed and its mean (see Fig. 3c). The base accuracy decreases when the environment gets lower in mean or slower in time scale. In particular, a slow controller achieves worse accuracy in a slow environment compared to a fast one. As a contrary effect, as the controller speeds up, it departs to a better accuracy later in a fast environment than in a slow one. That is why in Figure 3c the accuracy curves for slow and fast environment intersect, which also explains the local maxima in 3b. The slower environment - although having a lower base accuracy as a handicap - can be compensated for (in terms of accuracy) already at a slower controller speed.

**Fig. 3.**
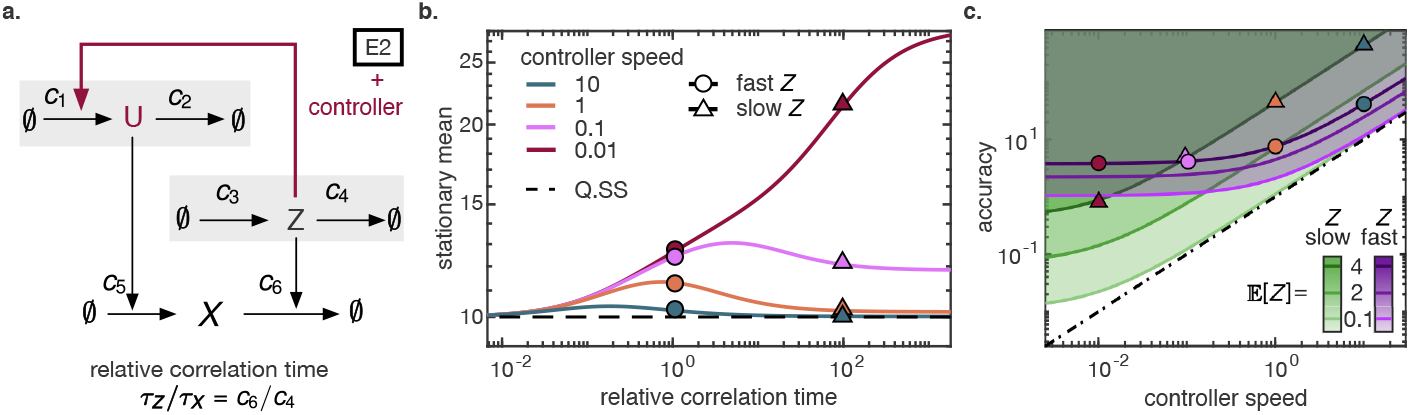
**a.** Reaction scheme for the environment with controller. The birth-death environment E2 is mitigated by a controller that senses the environment and regulates the birth rate of *X* accordingly. **b.** The stationary mean as a function of the relative correlation time *τ_Z_/τ_X_*. Different relative controller speeds *c*_2_/*c*_6_ are plotted. Circles (triangles) indicate fast (slow) environment. **c.** Accuracy Δ^−1^ as a function of controller speed *c*_2_/*c*_6_ for different values of environment mean 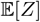 and speed *c*_4_/*c*_6_. Green (purple) indicates slow (fast) environment. The mean value 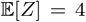 is highlighted by triangles (slow) or circles (fast) to match **b**. The slow environment achieves lower accuracy than the fast one for inactive controller (controller speed near 0). In order to improve accuracy, less increase in controller speed is needed for slow than for fast environment. As a consequence the accuracy curves for slow and fast environment intersect. The dashed line indicates the the critical controller speed needed to reach a given accuracy level for all environment mean and speed. The slow and low environment exhausts this universal accuracy-controller relation. Parameters were 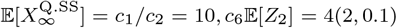, *c*_6_/*c*_4_ = 1 (circles, purple), 100 (triangles, green).

As a key question, we asked at which time scale the controller must operate to mitigate environmental perturbations with a given accuracy. To formalize this, we request the deviation in Eq. (25) to stay below a critical margin Δ* or the accuracy to stay above 1/Δ*. Interestingly, we found that for each accuracy margin, a critical controller speed can be chosen that operates universally for all environment speeds and means (see the dashed line in Fig. 3c). For fixed *c*_5_/*c*_6_ and *c*_1_/*c*_2_, the critical relative controller speed is given by

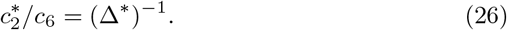

When the environment changes in mean or time scale, the controller holds the accuracy within the tolerated margin. Since the robustness that we analyze is defined via the steady state behaviour, our statement is restricted to environmental changes that occur so rarely that the steady state can be reached between changes (see Appendix Fig. B4).

Which environment speed and mean exhaust the critical accuracy? It is not the fast and furious (i.e., large mean) environment that causes deviation from Q.SS. For fast environment, the degradation rate of *X* averages out to the mean, making the Q.SS assumption valid. For large (furious) environment, the main driver of deviation 0 is hardly visited. In addition, the furious environment boils up the controller to act in a regime where it has a higher signal-to-noise ratio. This leaves the slow and low *Z* to exhaust the critical deviation. During excursions (*Z* = 0), the average dynamics follows

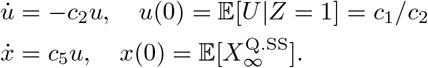

Consequently, the deviation for an infinitely long excursion rises, on average, to the value

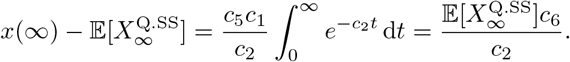

The excursion becomes the single dominating share as 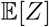 tends to 0, and *τ_Z_* → ∞ justifies infinitely long excursions. Since this dominating case is linear, the average dynamics is justified and the heuristic derivation of Eq. (26) can be made rigorous.

In summary, we observed that the accuracy margin was exhausted by the regime of slow and low environment, leaving the controller with the simple task to react as fast as to guarantee that a plateau is reached within the tolerated deviation, on average. This finding joins the variations on the theme ‘faster sensor molecules achieve higher accuracy’. Note that the setting and the control objective differ from [48]. In the latter, a sensor molecule recorded the progression of the target species, while here it senses the environment. The objective of suppressing fluctuations, i.e., the variance, in the controlled species induced the quartic root law on the sensing event counts. Here, in contrast, we asked for the stationary mean to stay within a tolerance of the setpoint, and this induced an inverse proportional law on the speed with which the controller responds to the environment. To summarize, both findings show that, when under the influence of a random environment, the accuracy can be increased at the cost of a faster sensor molecule. Our finding stands out due to its independence of size and speed of *Z* and the proportional relation in Eq. (26).

### 4.3 Stochastic toggle switch in a random environment

In the previous examples, we observed the effects of the modulation of the stochastic birth-death process given different architectures of this process. The next question of interest is whether the modulation effects propagate into the networks constructed from several such processes that interact with each other. Here, we consider a simple genetic toggle switch without cooperative binding, which is one of the best-studied small gene networks that induces bistability [49]. A balance of a toggle switch is known to be sensitive to noise [50], and we hypothesise that the effect of the noise in the stochastic environment should be visible at the level of the mean expression of the component genes.

We model the genetic toggle switch as a subsystem of two processes that are modulated by a shared environment (*Z*_1_, *Z*_2_) in the same way as the single gene is modulated in the section 4.1.2. These processes interact with each other by reducing the birth rate of their counterparts (Fig. 4a). In terms of a gene network, each process here is the number of proteins expressed by a gene. Their birth and death rates are modulated by the environment, and the birth rates are decreased in the presence of proteins expressed by the other gene. This model can be described by the following set of stochastic reactions:

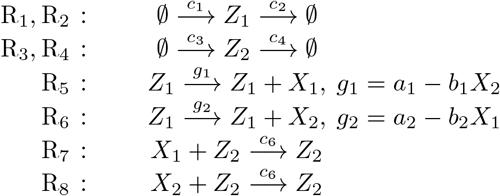

**Fig. 4.**
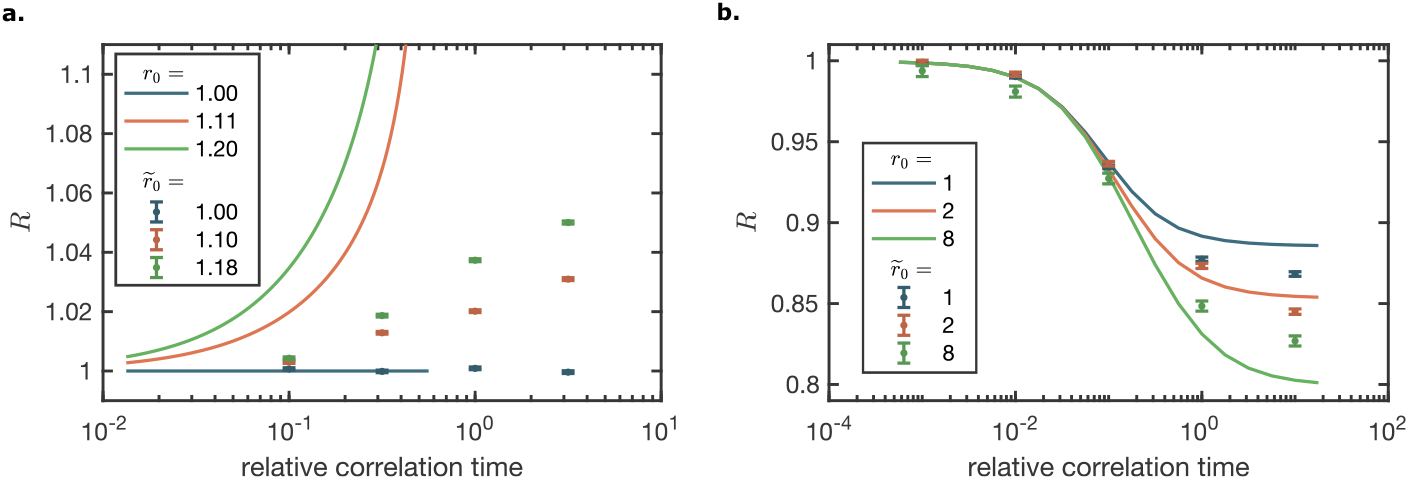
The relative asymmetry change ratio *R* and its counterpart 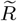 as a function of the relative correlation time *τ_Z_/τ_X_* for different baseline asymmetry *r*_0_ and 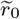 values, computed for the subsystem species of **a.** toggle switch with fixed parameters 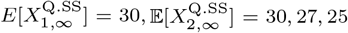, *b*_1_ = *b*_2_ = 0.02, *c*_6_ = 0.01, with *a*_1_, *a*_2_ matched accordingly; and **b.** oscillator with fixed parameters 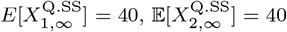, *b*_1_ = 0.02, *c*_6_ = 0.01, with *a*_1_ and *b*_2_ matched accordingly. The fixed parameters of the environment of both toggle switch and oscillator are: *c*_1_ = 300, *c*_2_ = 100, 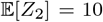. All simulations were performed for 10^6^ time points.

Reactions R_1_, R_2_, R_3_, R_4_ model the stochastic environment. Reactions R_5_, R_6_ describe gene expression and regulation modulated by the stochastic environment *Z*_1_. Rates *a*_1_, *a*_2_ correspond to the rate of expression of a the fully active gene, whereas coefficients *b*_1_, *b*_2_ are proportional to the strength of repression of the corresponding regulatory molecules produced by the counterpart gene. Finally, reactions R_7_, R_8_ describe the decay of the gene products modulated by the stochastic environment *Z*_2_. The subsystem species *X*_1_ and *X*_2_, besides interacting directly, are additionally coupled by the confounding factors *Z*_1_ and *Z*_2_.

Note that here we use a linear function to model repression, instead of, e.g., a Hill function, which would be a standard choice for this purpose [51, 52]. This decision is based on the fact that a standard repression model employs a non-linear reaction propensity, whereas ESME can work only with linear subsystems. In order to adhere to an implicit assumption that numbers of each species of the subsystem cannot be negative, we only considered the region of the parameter space of the model where the following is true. First, no share contributed by any state of the environment can be negative. Second, the total share contributed by the states that stabilize at negative values of the subsystem species cannot exceed 0.01%, which keeps its impact on the modulated mean expectation negligibly small (see also Appendix C.1).

To investigate the effect of the environmental modulation on the asymmetry of the toggle switch, we first define the asymmetry of the switch as a ratio of the stationary means of its component processes:

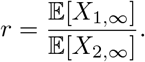

In the case of a constant environment, we denote this ratio as *r*_0_, the baseline asymmetry of the switch:

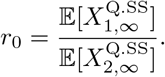

Then we can quantify the effect of a given environmental modulation on this asymmetry by computing the relative asymmetry change ratio 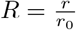.

Here, we chose the model parameters that result in three different *r*_0_ ratios and, for each of these cases, varied the relative correlation time 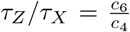 while keeping the ratio 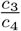 fixed. The stationary means for various *τ_Z_/τ_X_* were computed with ESME (Appendix B.6). From Figure 4a (solid lines), (i) a symmetric switch stays symmetric also in a random environment, (ii) the relative asymmetry in an originally asymmetric switch increases with the relative correlation time, (iii) this effect of the environmental speed on the asymmetry is higher when the baseline asymmetry, *r*_0_, is higher. We also note that, for asymmetric switches, the value of *R* grows towards a vertical asymptote. This asymptote signifies the *τ_Z_/τ_X_* value at which the weaker gene of the toggle switch (in our example, it is *X*_2_) becomes repressed so strongly that its mean approaches 0. The switch model with the repression implemented using linear propensities is descriptive of the modelled system only on the left side of this asymptote (for limitations, see Appendix C.1).

Given that the asymptote is an artifact of the linearization in reactions R_5_, R_6_, we verified that, apart from it, the observed monotonic dependencies (i), (ii), (iii) of the relative switch asymmetry are indeed the property of the toggle switch in a random environment. For this, we replaced the R_5_ and R_6_ with reactions 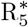 and 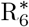

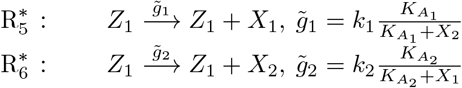

that utilize a Hill function (without cooperative binding), which is a standard way to describe the expression of a gene subject to repression [51, 52]. Since the exact stationary mean is computationally unfeasible to obtain, we performed stochastic simulations [14, 53] of the changed model. To ensure that the models match, we performed a ‘‘reverse linearization” between the linear and the Hill function repression models (Appendix C). Furthermore, we chose parameters of the reactions 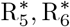 such that *r*_0_ and 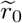 take similar values for convenient comparison.

From Figure 4a, we see that all the three observations (i), (ii), (iii) made with regards to how *τ_Z_/τ_X_* affects the asymmetry of a toggle switch hold for the model with a nonlinear repression function. However, while the behaviour is the same in terms of the symmetric case (i) and the monotonic properties (ii), (iii), there is a shift along the x-axis between the *R* and 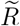 values. This is likely because the Hill function represses less strongly than its linearization. For the linearized repression, the dominant stable state with high *X*_1_ and low *X*_2_ is induced at a lower perturbation, i.e., at lower correlation, even driving the *X*_2_ to full extinction. We conjecture, that an asymptote also occurs for the Hill model, caused by much higher *X*_1_ values than *X*_2_ values due to excursions, likely also driving *X*_2_ to full extinction.

### 4.4 Stochastic oscillator in a random environment

While some effects of the random environment on the mean levels of the toggle switch components were to be expected, it might seem counter-intuitive that an effect will also be visible in a network with a periodic behavior. In particular, one might object that, in an oscillatory system, the information about the network period is lost when we move from the time series to the mean levels in the analysis. However, an oscillator is also characterized by an amplitude of its components, and we expect that the effects of the random environment on these amplitudes will be visible in the mean levels of the subsystem components. Further, changes in the dynamics of skipped oscillations, which are possible in stochastic oscillators, could propagate into changes in the mean species levels.

Reactions of the oscillator model differ from the reactions of the switch model (section 4.3) only in reaction R_6_, which now models induction of the species *X*_2_ by the species *X*_1_ with the induction strength coefficient *b*_2_ and takes form:

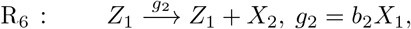

Note that here, as in section 4.3, we model repression propensity with a linear function. Thus, we investigated the model only in the subset of its parameter space where no share contributed by any state of the environment is negative and the mean values of the subsystem species stabilize at non-negative values in all considered states of the environment.

We quantified the effect of a given environmental modulation on this oscillator by computing the relative asymmetry change ratio 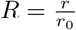, as in section 4.3, in three different *r*_0_ ratios over the varied relative correlation time 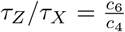 while keeping the ratio 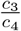 fixed. The stationary means for various *τ_Z_/τ_X_* were computed with ESME (Appendix B.7). From Figure 4b (solid lines), *R* decreases with the relative correlation time for all *r*_0_ values, and this effect is more pronounced at higher *r*_0_ values. In terms of *X*_1_ and *X*_2_, this means that a random environment causes 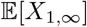 to decrease relative to 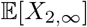, and this effect is stronger when the oscillator has higher 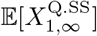 relative to 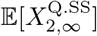.

As in section 4.3, we verified that the observed effects are the property of the studied network in a random environment and do not qualitatively depend on the linearization assumption made when modelling repression in reaction R_5_ by performing stochastic simulations of the oscillator model with R_5_ replaced by 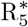. From Figure 4b (dots), the results of the simulations are qualitatively the same as those produced by ESME.

## 5 Discussion

In this work, we provided an exact stationary mean evaluation for linear chemical reaction networks in Markov environment, as a way to address (i) the difficulty of deriving the means in networks with bimolecular reactions and (ii) the computational infeasiblity of co-simulating the environment with the subsystem. Namely, the computation of stationary means in chemical reaction networks commonly relies on Monte Carlo simulation, moment closure or the approximation via a linear program with convex constraints. These approaches are often time-consuming or prone to approximation errors. Our expression computes the stationary mean exactly for a particular class of chemical reaction networks, i.e., the ones that can be decomposed into a linear subsystem and an environment, where the modulation is only allowed unidirectionally from environment to subsystem. Often such decomposition does not exist, because the requirements of linearity and unidirectional modulation are limiting. Common violations are the following: (1) the existence of a reaction *A* + *B* → *C* where at least one of the three species is in the subsystem, e.g., the MAPK/ERK pathway [54]. If *A* is an environmental species, it needs to be preserved during the reaction, and *A* and *B* cannot be both in the subsystem due to linearity. (2) Systems for which any partition into two sets of species comprises bimolecular reactions in both directions are not feasible. Finally, (3) reaction rates that have a Hill or other non-linear dependency violate the linearity constraint. Additionally, our expressions require the stationary distribution of the environment, which further limits our approach.

While models on continuous state spaces, e.g., Langevin equations, can often handle rates that become temporarily negative, we found that our method can be sensitive to this situation when summands in Eq. (10) get negative. This can prohibit rate linearization. When we compute ESME numerically, we need a truncation of the environment to a finite state space. For this, the finite state projection method [22, 55] or other tools [56] can be employed.

In the experimental context, we suggest that the effect of an environmental embedding can be distinguished from the Q.SS model with averaged environment via a characteristic fingerprint at the bulk level. Single stationary mean values are hardly informative to distinguish between both. However, functional dependencies over system parameters can look qualitatively different for a Q.SS model compared to an extension via an environmental embedding. In this work, we mostly investigated the time scale separation as a system parameter and saw a characteristic qualitative deviation. While the Q.SS model remained constant, the environment model generated a dependency with phases of distinct functional behaviour. It is hard to modify the time scale separation experimentally. However, other system parameters, such as volume, temperature, inducer concentration or time, can yield dependencies that achieve a phase change. In this way, the effect of the environment could be seen at the bulk level.

Reduction techniques for the stochastic simulation of chemical reaction networks can have difficulties in capturing the asymptotic mean correctly [57, p.70]. Our approach might assist in detecting the parameter regimes in which approximate simulations succeed. On the one hand, ESME can replace the Monte Carlo approximations. On the other hand, our quantification of environmental shares can permit insight into the failure mode of the approximate simulation or model reduction technique. The knowledge of the exact stationary mean can also tune approximation methods towards capturing the asymptotic mean correctly.

In future work, we seek to derive expressions for the stationary variance and generalize the expressions to a semi-Markov environment, analogously to [39], and to environments on continuous state space.

## 6 Statements and Declarations

Partial financial support was received from Marie Skłodowska-Curie Actions Fellowship (grant agreement ID: 101033300 to Sofia Startceva). The authors declare no competing interests.

## Appendix A Proofs of Proposition 1 and the main result (ESME)

### A.1 Proof of Proposition 1

*Proof* We verify the stationarity condition for the embedded chain.

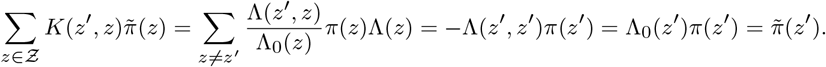

### A.2 Proofs of proposition 2 and theorem 3 (ESME)

In the proofs we make use of the following lemma.

#### Lemma 4

*Let T be exponentially distributed with parameter μ* > 0 *and suppose that all eigenvalues of* **A** *have non-negative real part. Then μ* **I** + **A** *is invertible and*

i. 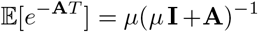
ii. 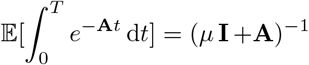
iii. 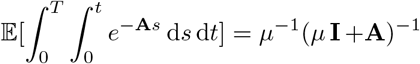

*Proof of the lemma*

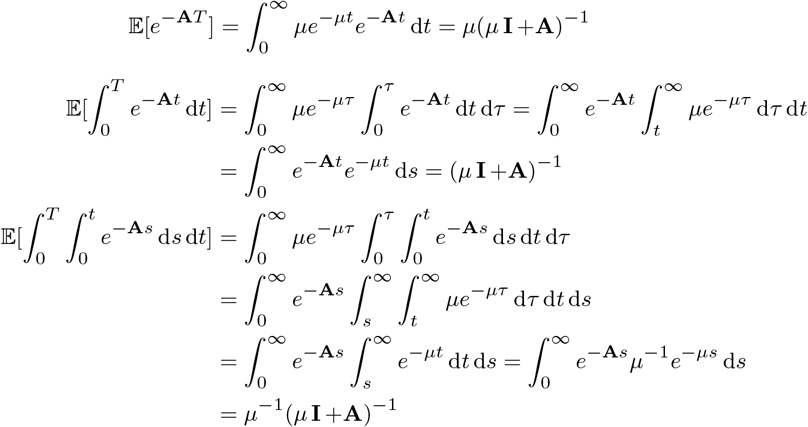

*Proof of proposition 2*

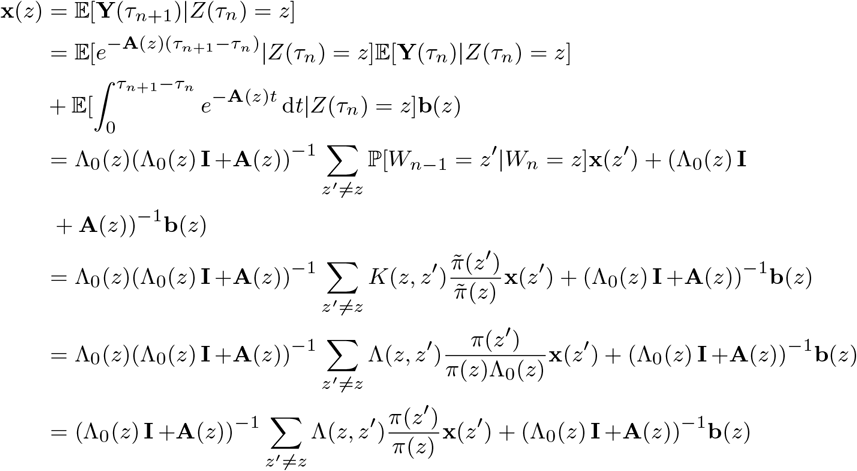

Hence,

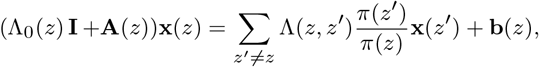

or upon adding of Λ(*z*, *z*)**x**(*z*):

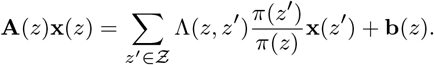

If detailed balance hold, we substitute

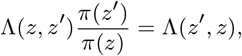

yielding

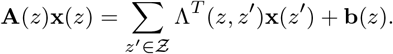

*Proof of* Eq. (17) The integrals evaluate to:

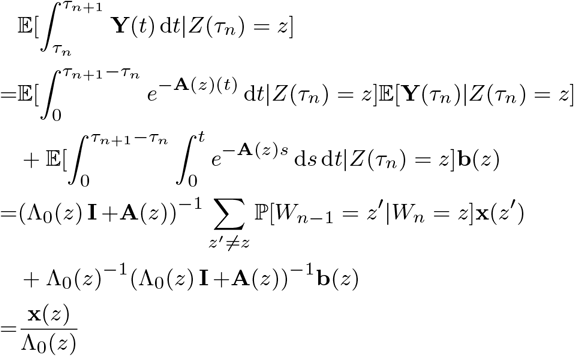

*Proof of theorem 3*

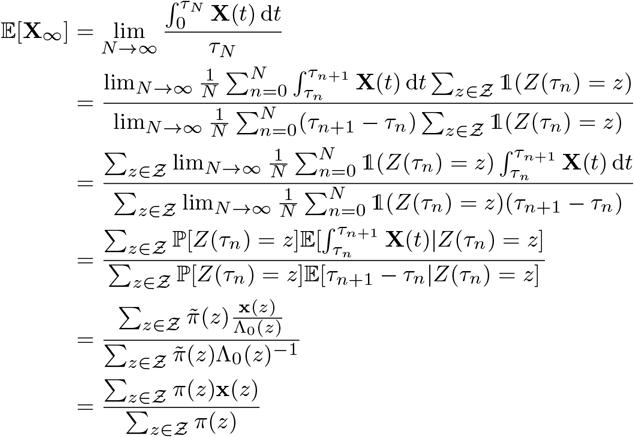

## Appendix B Details on Case Studies

### B.1 E1: Two-state death rate modulation

The birth rate is a birth-death process and the death rate is a two-state Markov process (random telegraph model), i.e., consider the CRN (see Fig. 2a.)

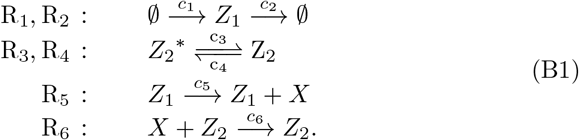

The stationary means of *Z*_1_ and *Z*_2_ are easily identified as

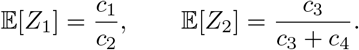

In this case, the dimension of the subsystem is *d* = 1. Then *A*(*z*_1_, *z*_2_) = *c*_6_*z*_2_ and *b*(*z*_1_, *z*_2_) = *c*_5_*z*_1_ are scalars. Since *Z*_1_ only enters via the zero-order reactions, i.e., via b, we use the Q.SS assumption and set it to its mean. This reduces the environment to *Z* = *Z*_2_ on the state space 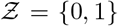. The generator of the environment is

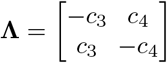

with stationary distribution *π* = Bernoulli(*c*_3_/(*c*_3_ + *c*_4_)). The expression of the stationary mean is then (Eq. (12))

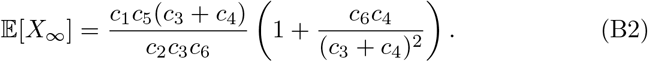

We analyzed the behaviour of the stationary mean 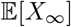 as a function of the relative correlation time *c*_6_/(*c*_3_ + *c*_4_).

### B.2 E2: Birth-death modulator

In Eq. (B1), replace the two reactions R_3_, R_4_ by

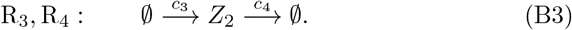

The full CRN is visualized in figure 2b.

While the collection of rates **A**, **b** of the *X* dynamics remains the same, we make adjustments in the stationary mean state space, generator and stationary distribution of *Z*_2_ accordingly:

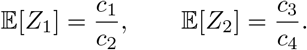

The standard birth-death generator on state space 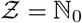 is given by

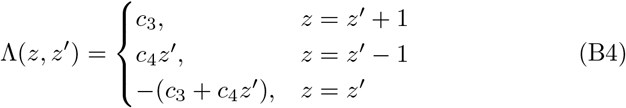

with stationary distribution *π* = Poisson(*c*_3_/*c*_4_). The expression of the stationary mean was already presented in [39] and [37]. Analogously to model E1, we analyzed the behaviour of the stationary mean 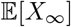 as a function of the relative correlation time *c*_6_/*c*_4_. For computational purposes, we truncate the state space 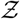 to 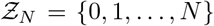 with a large enough *N*. Here, *N* = 99 was used.

**Fig. B1.**
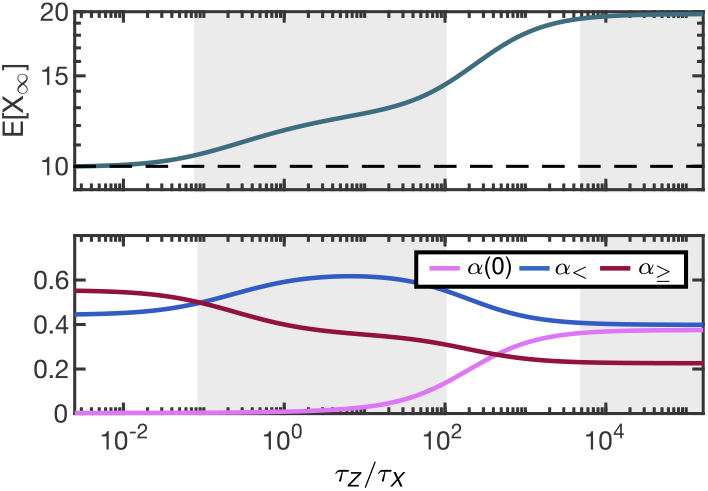
Model E2 with leakage. **Upper panel**. Stationary mean as a function of ln(*c*_4_). The stationary mean systematically exceeds the Q.SS mean 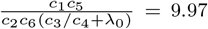. Four regimes can be distinguished: the asymptotic regime, the intermediate regime, the rising degenerate regime and the saturating degenerate regime. **Lower panel**. The contributions of *Z*_2_ to the stationary mean. Parameters were 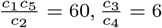, *c*_6_ = 1, λ_0_ = 0.02.

**Fig. B2.**
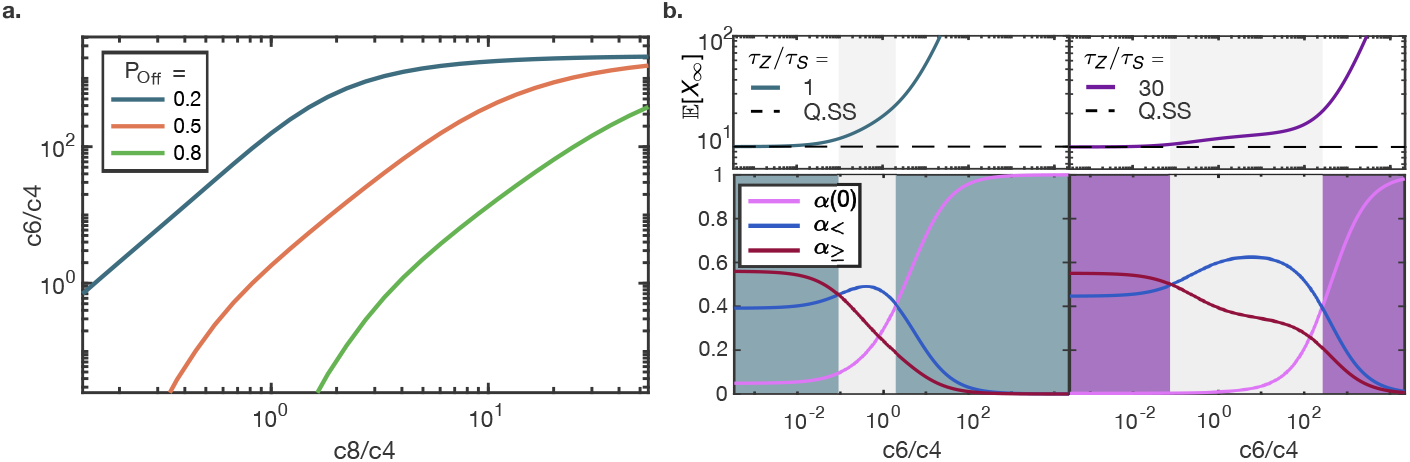
Model E3 Mutable *c*_3_ synthesis. **a.** The figure shows the *c*_4_ value at which *α*(0) and 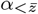 intersect as a function of 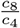. Dominant 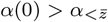 indicates that the stationary mean has entered the degenerate regime. The three curves illustrate different values of 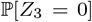. For 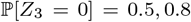 and slow enough relative speed *c*_8_/*c*_4_ no intersection was found, because *α*(0) dominated 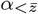 for all *c*_4_. The mean 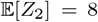 was fixed and *c*_3_/*c*_4_ adapted accordingly. Parameters were *c*_1_ = 0.4, *c*_2_ = 0.01, *c*_5_ = 1, *c*_6_ = 0.5 and 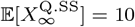. **b.** The left upper panel shows the slow switching *Z*_3_, whereas the right panel shows the fast switching *Z*_3_, compare Figure 2f. The corresponding shares are depicted in the lower panels. Parameters were as in Figure 2f.

#### B.2.1 Leakage

Compared to model E2, we include a base degradation rate λ_0_ for reaction R_6_, i.e., the propensity of reaction R_6_ is *f*(*x*, *z*) = (*c*_6_*z* + λ_0_)*x*. This altered **A**, i.e., **A**(*z*) = *c*_6_*z* + λ_0_. The figure B1 indicates four regimes, in particular the three phases of the model E2 analysis (Fig. 2e.) persist. The fourth phase with largest *τ_Z_/τ_X_* reaches saturation.

### B.3 E3: Mutable synthesis of the modulator

We modified reaction R_3_ of model E2, see Eq. (B1) and Eq. (B3), to

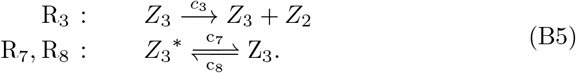

Then the two-dimensional environment 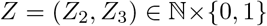 has a stationary distribution that is expressed via the confluent hypergeometric function. Expressions for 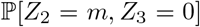 and 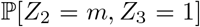 were derived analogously to [58] by expanding the generating function given therein. Define 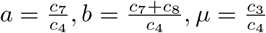. Then it holds

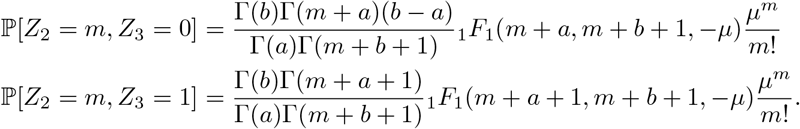

ESME was calculated numerically using Eq. (11) with truncation *N* = 100. Figure B2a shows the entry of the stationary mean into the degenerate regime as a function of the relative speed of the modulator 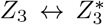. Figure B2b depicts the environmental shares for the slow and the fast modulator.

### B.4 Asymptotic behavior in a slow environment

Consider any of the models E1, E2, E3. Under the fixation of 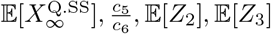 and *τ_Z_/τ_S_* the stationary mean only depends on the relative time scale *τ_Z_/τ_X_*. Hence in the following we fix 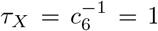. Then, *τ_Z_/τ_X_* → ∞ is equivalent to *c*_4_ → 0. By our main theorem (3), we obtain 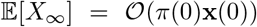 for *c*_4_ → 0 in the models 1a, 1b and 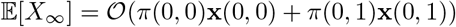 for model E3. By definition 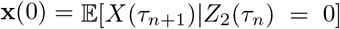. The function 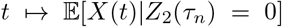 progresses affine-linearly in time. For very slow *Z*_2_ the excursions are long and the base value at *τ_n_*, from which they depart, is small compared to the value they reach at *τ*_*n*+1_. We thus neglect the base value to obtain a linear progression with constant slope 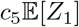, yielding

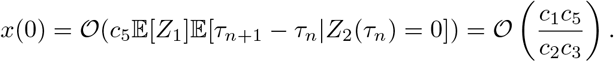

In total we obtain for the stationary mean of models E1, E2

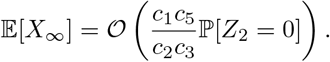

Since 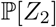 only depends on the mean 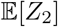 which we keep fixed, the parameter *c*_3_ which scaled linearly with *c*_4_ dominates the asymptotic behaviour *c*_4_ → 0. More precisely, we obtain

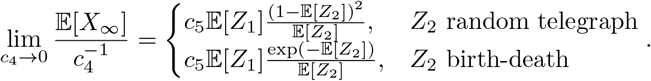

For model E3 the two states (*Z*_2_, *Z*_3_) = (0, 0) and (0, 1) contribute for *c*_4_ → 0. After setting up the recursion that couples *x*(0, 0) and *x*(0, 1), see next paragraph, we obtain

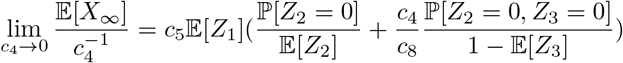

Note that 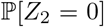 and 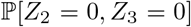 only depend on the relative rates 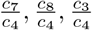, which were fixed, i.e., the time scale of the joint environment (*Z*_2_, *Z*_3_) was varied. The inverse proportional dependence on *c*_4_ for models E1 - E3 is visualized in Figures 2c.,f.,i.

For *c*_4_ → 0 we compute the stationary mean under assumption that all terms in Eq. (10) vanish except for *z* = (0, 0), (0, 1). The recursion equations (7) for the two states (0, 0) and (0, 1) read

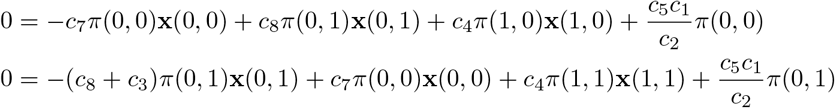

By assumption,

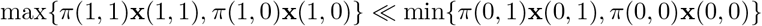

so we can set *π*(1, 1)**x**(1, 1) = *π*(1, 0)**x**(1, 0) = 0. Then the 2 dimensional linear system has the solution

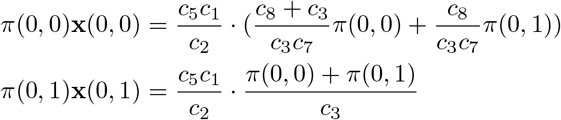

which by 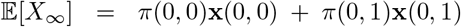 yields the result for 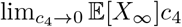.

**Fig. B3.**
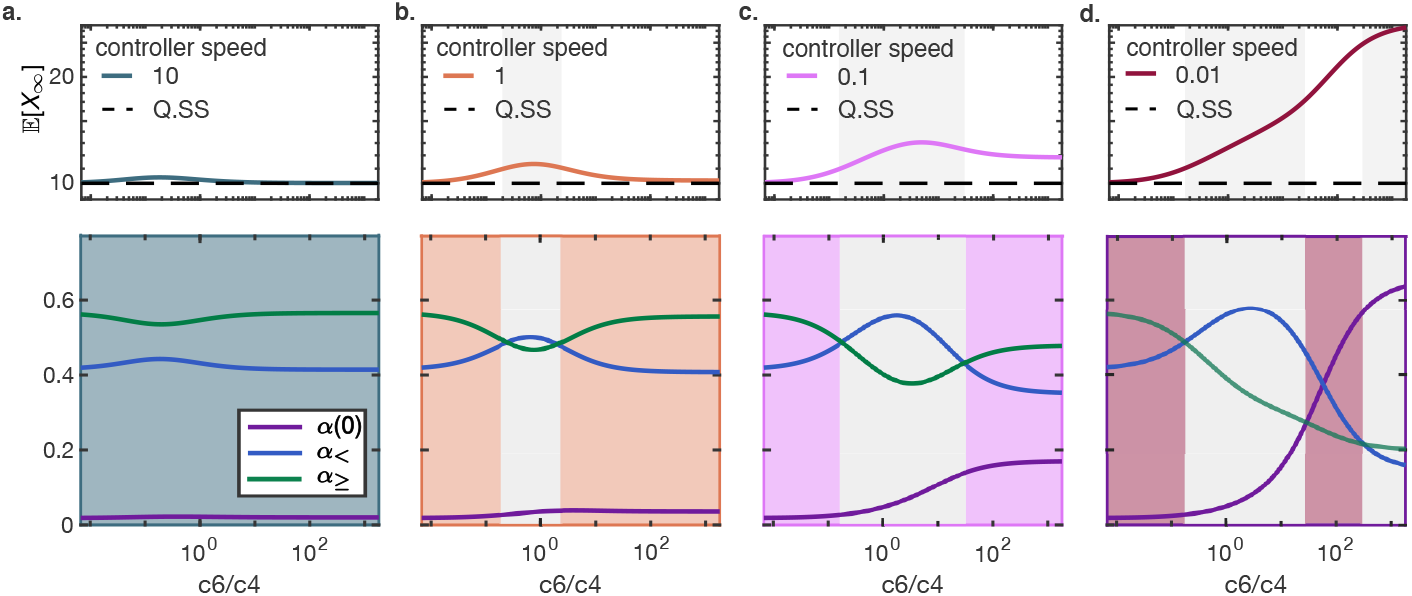
The controller *U* becomes slower from **a** to **d**. Parameters were *c*_5_ = 1, *c*_6_ = 1, *c*_1_/*c*_2_ = 10, *c*_3_/*c*_4_ = 4. For small *c*_6_/*c*_4_ the stationary mean saturates to the Q.SS mean *c*_1_ *c*_5_/*c*_2_*c*_6_ = 10 independent. The saturation level for large *c*_6_/*c*_4_ depends on the speed *c*_2_ of *U*. For fast enough *U* a local maximum appears. For slower *U* the saturation level increases and the contribution *α*(0) dominates more and more.

### B.5 E2 + controller: a correlated environment

Compared to environment E2, we replaced the reaction R_1_ in Eq. (B1), (B3) by

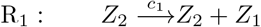

In order to apply our general framework we regard *Z*_2_ as the one-dimensional environment and (*Z*_1_, *X*) as the modulated linear CRN. Then 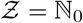 and Λ is the generator of a birth-death process with birth rate *c*_3_ and death rate *c*_4_. Detailed balance is satisfied by Poisson-distributed π with parameter *c*_3_/*c*_4_. For the dynamics of **Y** we obtain

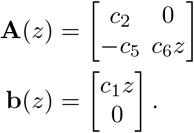

We fix *c*_6_, *c*_5_ and the means 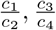. For different speeds *c*_2_ of *Z*_1_ we let the speed *c*_1_ of *Z*_2_ vary, see Figure 2k. For large *c*_4_ the stationary mean saturates to the Q.SS mean *c*_1_*c*_5_/*c*_2_*c*_6_ = 10 independent of *c*_2_. The saturation level for small *c*_4_ depends on the speed *c*_2_ of *Z*_1_. For fast enough *Z*_1_ a local maximum appears. For slower *Z*_1_ the saturation level increases. The stationary mean curve for small c_2_ resembles the leakage case displaying the same four phases.

The birth-death process *X* in a correlated environment showed a local maximum in the phase analysis. Here we provide details on the shares 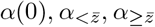, see Figure B3. Furthermore, Figure B4 provides example trajectories for a fast and a slow controller.

**Fig. B4.**
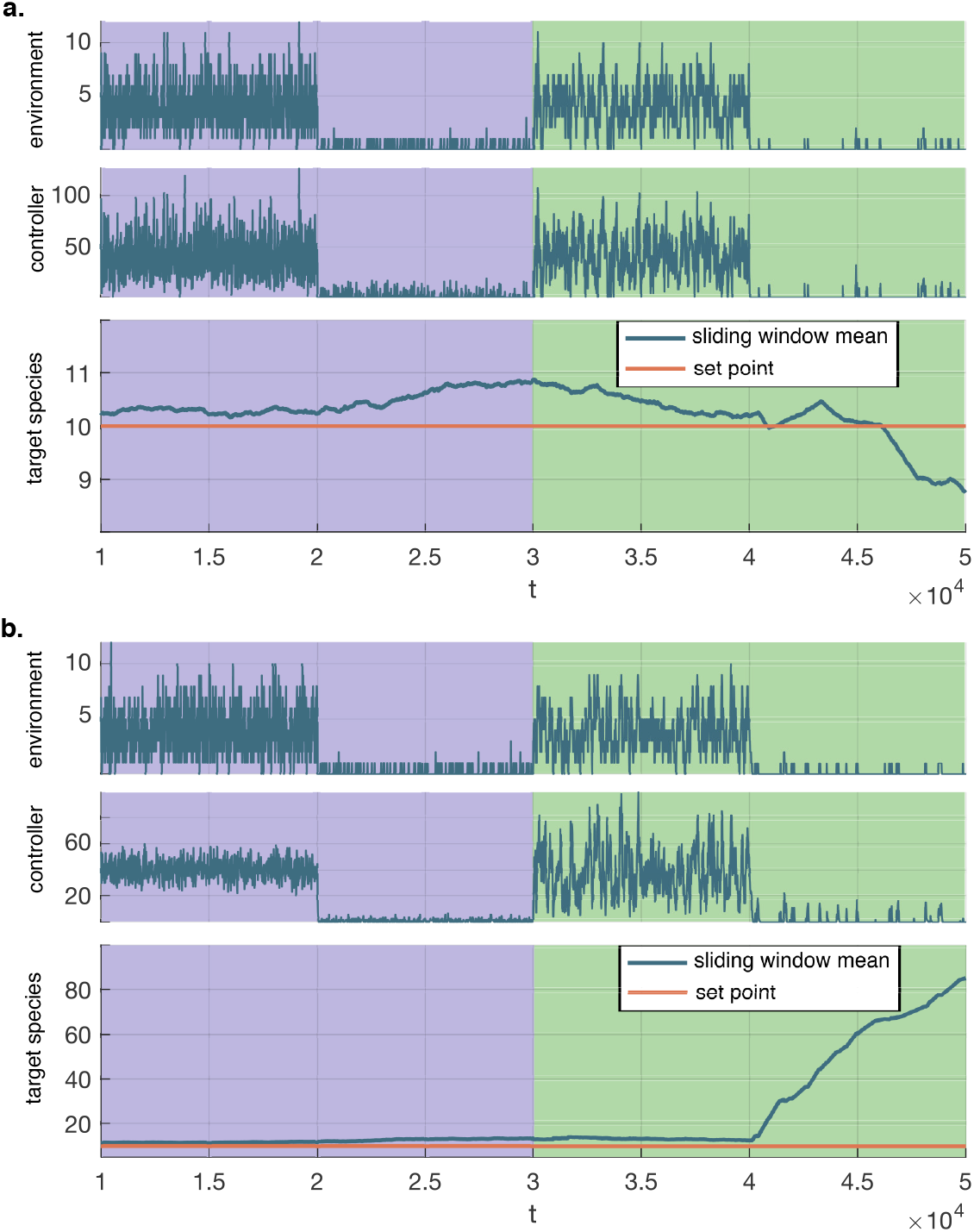
Trajectories for the environment E2 with controller. Environment *Z* and controller *U* are shown in plain form, while *X* is depicted as a moving average with window length *t* = 10, 000 to match the statement about the stationary mean. A burn-in time of *t* = 10, 000 was used. The environment progresses through four stages of different mean and speed. **a**. A fast controller (*c*_2_/*c*_6_ = 10) keeps the target species near the setpoint, on average, as the environment changes. According to the theory the deviation stays within 10% of the setpoint. In the last phase a higher deviation indicates that stationarity is not reached within the window length. The drop below the setpoint is due to a low plateau value in the trajectory that would be counter-balanced by larger plateau values in the long run. **b.** A slow controller (*c*_2_/*c*_6_ = 0.1) achieves low accuracy for the critical stage of low and slow environment (last phase). Parameter values were 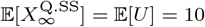, *c*_6_ = 1 and in the four phases the four pairs 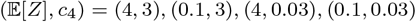 were used.

### B.6 Stochastic toggle switch in a random environment

Given the model reactions, the dynamics of 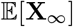 can be described with the following reaction matrices:

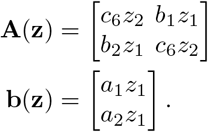

The generator used here is the one corresponding to two jointly independent birth-death processes, and it is of the form

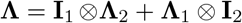

with 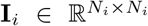 the identity matrices for state space truncation *Z_i_* ∈ {0,1,…, *N_i_* – 1} and **Λ**_*i*_ analogous to Eq. (B4). Here, *N*_1_ = *N*_2_ = 50 was used.

To pick the model parameters for a given *r*_0_ ratio, we fixed all parameters besides the rates *a*_1_ and *a*_2_ and computed those rates from the equation 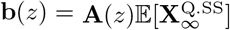.

### B.7 Stochastic oscillator in a random environment

Given the model reactions, the dynamics of 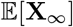 can be described with the following reaction matrices:

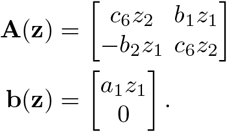

The generator used here is the one corresponding to two jointly independent birth-death processes, and it is of the form

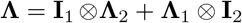

with 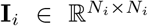 the identity matrices for state space truncation *Z_i_* ∈ {0,1,…, *N_i_* – 1} and **Λ**_*i*_ analogous to Eq. (B4). Here, *N*_1_ = *N*_2_ = 50 was used.

## Appendix C Details on linearization

We matched the linearized and the Hill function models of repression as follows. Namely, first in *g*_1_, we obtained

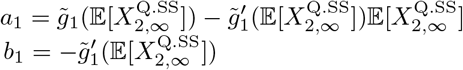

and then in 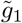 we expressed *k*_1_ and *K*_*A*_1__ in terms of *a*_1_ and *b*_1_ (analogously, the same is done for *k*_2_ and *K*_*A*_2__):

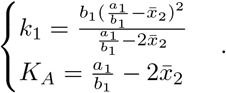

### C.1 Limitiations

Here, we elaborate on our understanding of the limitations of the linearization in the switch and oscillator case studies.

- First Limitation: As soon as *X*_1_ or *X*_2_ get too large, the propensities of the reactions R_5_, R_6_ become negative. This can happen for any choice of non-negative rate constants. Consequently, the linear dynamics Eq. (6) conditioned on *Z* = *z* can saturate at a state with a negative value for a subsystem component. This can, for large enough waiting time, cause the integral in the share Eq. (18) to become negative, and can be detected by checking for negative shares. Eventually, this can result in a negative stationary mean. In the toggle switch case study we checked that in the portrayed parameter range (i) no share is negative and that (ii) the total share of those environmental states for which the saturation point has a negative component, is sufficiently small.
- Second Limitation: Negative eigenvalues of the matrix **A**(*z*) can arise for some values of *z*. This means that the linear dynamics Eq. (6) conditioned on *Z* = *z* gets unstable. If the waiting time in the state is short enough or, equivalently, Λ_0_(*z*) is large enough, then the unstable dynamics is mitigated and the formulas in lemma 4 remain valid. This was not a limitation in the toggle switch case study due to a large enough *c*_1_.
- Third Limitation: The matrix **A** – Λ ⊗ **I** in Eq. (11) can get singular. We observed this in the toggle switch example for the choice *c*_4_ = 0.015 and other parameters that determine **A** – Λ⊗**I** as in Figure 4, causing an asymptote in 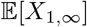 and 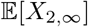. This parameter choice was, however, already excluded by the first limitation.

